# Glycochenodeoxycholic acid and ceramide suppress the antiviral effect of 25-hydroxycholesterol against human norovirus infection in human intestinal enteroids

**DOI:** 10.1101/2023.06.12.544665

**Authors:** Wadzanai P. Mboko, Preeti Chhabra, Anna Montmayeur, Ruijie Xu, Verónica Costantini, Jan Vinjé

## Abstract

The human intestinal enteroid (HIE) cell culture system with the support of glycine-conjugated bile acid glycochenodeoxycholic acid (GCDCA) and ceramide (C2) facilitate successful replication of several norovirus strains. Here we investigate how the presence of GCDCA/C2 impacts gene expression of norovirus-infected HIE and the impact of 25 hydroxycholesterol (25-HC), a key regulator of cholesterol homeostasis and bile acid production on norovirus replication. In absence of GCDCA/C2, 0.01 and 0.1 μM 25-HC suppressed virus (GII.4 Sydney[P16]) replication by 1.3 log and 1.1 log respectively (p<0.05). In the presence of GCDCA/C2, 5 μM 25-HC was required to achieve a 1 log decrease (p<0.05) in viral titers demonstrating that 25-HC restricts norovirus replication in HIE. RNA sequence analysis showed that during human norovirus infection, 25-HC downregulated expression of genes (CYP3A4, APOB, APOA1, and ABCG1) involved in cholesterol metabolism and transport as well as interferon stimulated genes such as ISG15 and IFIT1. GCDCA/C2 counteracts the suppressive effect of 25-HC expression of some genes related to these pathways including APOA4 and CYP27A1 however, other cholesterol genes such as APOA1 were further suppressed in the presence of GCDCA/C2.

**Importance:** Norovirus is the leading cause of epidemic and endemic acute gastroenteritis worldwide and currently, there are no effective therapeutic strategies against this highly contagious pathogen. Our study provides insights into the effect of bile during norovirus infection, highlight the role of the cholesterol/oxysterol pathways during human norovirus replication, and demonstrate the potential utility of oxysterols in developing norovirus therapeutics.

## Introduction

Human noroviruses are a leading cause of epidemic and endemic acute gastroenteritis worldwide (1–4). Currently, there are no licensed vaccines or antiviral treatments for this highly contagious pathogen although several vaccines are in various stages of development (5–7). The immune response to human norovirus infection remains poorly understood; however, the recently developed human intestinal enteroids (HIE) system has allowed to provide insights on the host factors that restrict virus replication (8, 9). HIE are primary cells derived from intestinal biopsies, maintained in a 3D culture, and seeded as monolayers to support human norovirus replication (10, 11). This novel cell culture system has revolutionized the norovirus field and enhanced our ability to study norovirus infection, assess efficacy of control measures, and measure adaptive and innate immune responses (12–14).

The interferon pathway is a key component of the innate immune response to virus infections and recent studies indicate type I and III interferon, the Jak/STAT pathway, and other signaling molecules (15–17) are pivotal in the control of human norovirus infection. Importantly, multiple other antiviral mechanisms may still be involved in restricting virus replication which may be important in developing strategies to mitigate norovirus related illness. Equally intriguing are the strain specific differences in sensitivity to interferon and requirements for bile acids during replication (10). Globally predominant GII.4 viruses can replicate independently of exogenous bile; however, virus replication can be enhanced by including bile acids in the cell culture media. Specifically, glycochenodeoxycholic acid (GCDCA) promotes GII.3 replication via a BA receptor sphingosine-1-phosphate receptor 2 (S1PR2) mediated mechanism, and enhances endosomal uptake and acidification, as well as activity of Acid Sphingomyelinase (ASM), an endosomal/lysosomal enzyme that generates ceramide on the apical membrane in HIEs (18, 19). How GCDCA affects the host response during human norovirus replication is an important question to address.

Over the last few years oxysterols have emerged as important regulators of innate and adaptive immunity (20). Specifically, 25 hydroxycholesterol (25-HC), an oxysterol that has been established as a key regulator of cholesterol homeostasis (21–23) and bile acid production (24) has now been demonstrated to have broad antiviral activity. Several studies have shown that 25-HC restricts replication of enveloped and non-enveloped viruses (25–30). A recent study also showed that pretreatment of RAW cells with 25-HC prior to infection with murine norovirus significantly decreased virus replication (31).

The interaction between 25-HC and human norovirus replication has also not been investigated. In this study, we aimed to determine the extent to which 25-HC restricts human norovirus replication in HIE and the interaction between 25-HC and GCDCA and ceramide (GCDCA/C2) during norovirus infection.

## Materials and Methods

### Human intestinal enteroid culture

Secretor positive jejunal 3D HIE cultures (J3 line) were cultured as previously described (8). After 7 days, 3D cultures were dissociated into a single cell suspension and plated as undifferentiated monolayers in 96-well plates. After 24 hours, differentiation media was added to induce monolayer differentiation and the plates were incubated for 4 days at 37°C. (Figure 1A).

**Figure 1:**
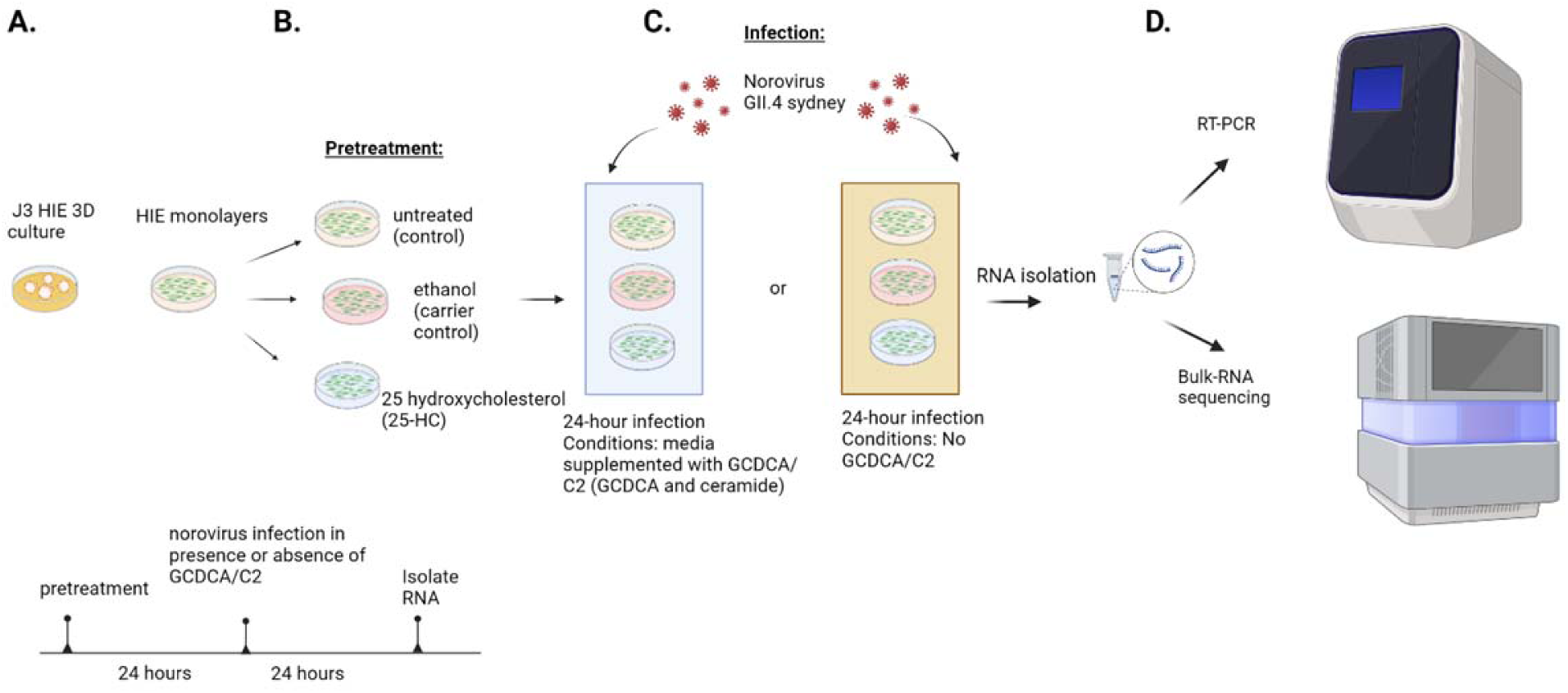
Experimental approach. A. J3 Human intestinal enteroid (HIE) monolayers were prepared from 3D cultures and differentiated for 5 days. B. HIE monolayers were treated for 24 hours with 24-HC, 25-HC, 27-HC, or the equivalent dilution of the carrier (ethanol) in media or untreated. C. After 24 hours of pretreatment, cells were infected with norovirus GII.4 Sydney[P31] in the presence or absence of GCDCA and ceramide (C2). D. Viral RNA was quantified using RT-qPCR and gene expression was determined by bulk RNA sequencing.

### Oxysterol pretreatment of HIE

Differentiated monolayers were treated with 0.01-5 µM 25 hydroxycholesterol (25-HC) as well as 0.1 µM - 2.5 µM 24 hydroxycholesterol (24-HC) and 27 hydroxycholesterol (27-HC) (Sigma-Aldrich). In parallel, cells were treated with equivalent concentrations of ethanol which was the carrier for the oxysterols, or differentiation media, as negative controls. Cells were treated for 24 hours at 37°C (Figure 1B).

### Infection and viral RNA detection

Pretreated monolayers were infected with a norovirus GII.4 Sydney[P16] positive 10% stool filtrate in infection media (CMGF-) for 1 hour at 37°C, then washed with CMGF-media and incubated for 24 hours in differentiation media. For infections performed in the presence of GCDCA/C2, infection media and differentiation media were supplemented with 500 μM GCDCA and 50 μM of ceramide (GCDCA/C2) (19). Cell lysates and media were collected at 1- and 24-hours post infection. RNA was extracted and viral RNA was detected by RT-qPCR (32, 33). (Figure 1C and D)

### Detection of cellular gene expression

RNA was prepared from cell lysates using the MagMAX™-96 Total RNA Isolation Kit (Thermofisher Scientific). RT-PCR was performed using the SuperScript™ III One-Step RT-PCR System with Platinum™ *Taq* DNA Polymerase and gene-specific primers (Table 1), or RNA was analyzed by bulk RNA sequencing (2×150 bp reads) on the Illumina NovaSeq (Figure 1D)

### RNA sequencing and data analysis

Raw sequencing reads were quality checked using FastQC v.0.11.5 (Babraham Bioinformatics - FastQC A Quality Control tool for High Throughput Sequence Data). Sequencing adaptors and low-quality reads were trimmed and filtered out using Fastp v.0.20.1 (34). Reads were pseudo-aligned to human reference transcriptome GRCH38.p13 with Gencode v39 annotations and with gene expressions quantified using the “quant” mode of Salmon v. 1.8.0 (35)

Genes differentially expressed between different experimental conditions were determined using the “DESeq” method in the “DESeq2” R package using the “Wald” test. Resulting p-values for each gene were then adjusted with the Benjamini-Hochberg method (36) to eliminate false positives (padj). Differentially expressed genes (DE genes) were defined as having an absolute log2fold change > 1 and padj value < 0.05.

Padj values were negatively log10 transformed and DE genes were visualized using the R package “ggplot2” (37). Relative expressions of select DE genes were further visualized using the R package “pheatmap”. The expressions of these genes were also clustered using the hierarchical clustering method to determine the relationships between samples.

### Pathway Enrichment analysis

Pathway enrichment analysis was performed with the ranked log2fold changes between RNA profiles of the two treatment groups included in the analysis with the “Proteins with Values/Ranks” function provided by STRING(38). Pathways enriched by the overlapped DE genes identified between 25-HC versus ethanol analyses performed with the presence and absence of GCDCA/C2 was performed with the “Multiple Proteins” function provided by STRING.

### Gene validation

Validation of RNA sequencing analysis for specific genes was performed by RT-PCR (Table 1). We selected genes that have been shown to be important in norovirus infection or upregulated by 25-HC treatment [14, 37, 38]. <u>Statistical analysis</u> Statistical analyses were performed using GraphPad Prism 8.0 software (GraphPad Software, La Jolla, CA). Treatment groups were compared by Mann-Whitney-test. Data are presented as mean ±SEM. All differences were considered statistically significant when the *p*-value was ≤0.05.

## Results

To determine the effect of GCDCA/C2 on HIE, we compared the gene expression profile of HIE monolayers incubated with and without GCDCA/C2 for 24 hours. We found 18 genes differentially expressed (DE) with log2fold change >1 in the HIE with GCDCA/C2 compared to the HIE without GCDA/C2. Immune defense related genes such as HLA-DPA1, surfactant protein A2 (SFTA2) and DDX24 were downregulated, while genes associated with intracellular regulation and transportation of lipids (ARHGAP33, ATP8B3, SCARF1) were upregulated (Figure 2A).

**Figure 2:**
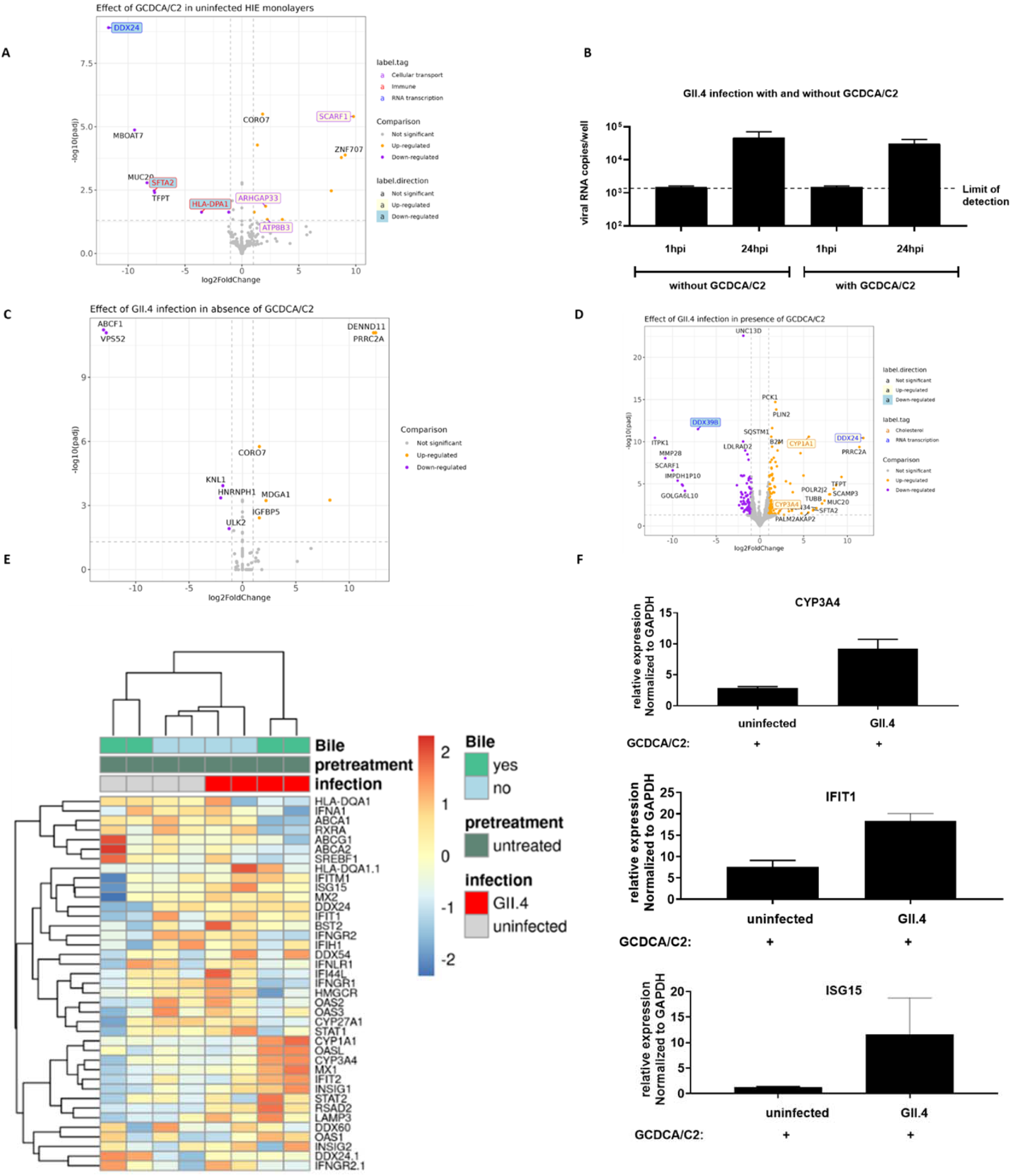
GCGCA/C2 affects cellular gene expression in infected and uninfected human intestinal enteroids. Differentiated HIE monolayers were infected with norovirus GII.4 Sydney[P31] in the presence or absence of 500 μM GCDCA and 50 μM of ceramide (C2). RNA was isolated and analyzed by qRT-PCR. A. untreated cells and cells treated with GCDCA/C2 for 24 hours RNA from uninfected cells or cells infected with norovirus was analyzed by bulk RNA sequencing to determine changes in cellular gene expression. Volcano plot for differentially expressed (DE) genes between: B. Quantification of norovirus RNA in cells infected in the absence of or presence of GCDCA/C2 at 24 hours post infection. Data represents summary of 3 independent experiments with 3-4 independent replicates per condition in each experiment., C. uninfected cells and cells infected with norovirus in the absence of GCDCA/C2, D. uninfected cells and cells infected with norovirus in the presence of GCDCA/C2. Genes upregulated by ethanol were colored in orange and genes downregulated by ethanol were colored in purple. E. Heat map depicting select genes. F. Normalized expression of CYP3A4, IFIT1, and ISG15 in HIE monolayers infected with norovirus GII.4 in the presence or absence of GCDCA/C2. RNA prepared at 24 hours post infection and analyzed by bulk RNA sequencing.

To study the effect of GCDCA/C2 on gene expression in HIE monolayers during norovirus infection, differentiated J3 HIE monolayers were infected with human norovirus GII.4 Sydney in the absence or presence of GCDCA/C2.There was no difference in viral titers in norovirus-infected HIE in the absence or presence of GCDCA/C2 after 24 hours of infection (Figure 2B). Transcriptionally, 11 genes were DE with log2fold change > 1 after GII.4 infection in the absence of GCDCA/C2 (Figure 2C) including ABCF1, an immune modulator (39, 40), which was downregulated in infected cells. In contrast, in HIE infected in the presence of GCDCA/C2, 206 genes were identified as DE. Genes associated with cholesterol metabolism (CYP1A1 and CYP3A4), and innate immune response to virus infection (DDX24) (41, 42) were upregulated after infection. Genes that facilitate RNA virus replication such as DDX39B [41], were downregulated (Figure 2D). GCDCA/C2 changes the magnitude of the transcriptional response in cells as depicted in the heat map (Figure 2E). Expression of select upregulated genes related to cholesterol and innate immunity pathways (CYP3A4, IFIT1 and ISG15) were confirmed by qPCR (Figure 2F).

Next, we sought to determine the extent to which GCDCA/C2 affects human norovirus titers and the cellular response in cells that were treated with 25-HC) prior to infection. First, we tested whether 25-HC restricts human norovirus replication by pretreating J3 HIE monolayers with a range of different concentrations (0.01-5 μM) of 25-HC in the absence or presence of GCDCA/C2. In the absence of GCDCA/C2, 0.1 μM and 0.01 μM 25-HC treatment reduced virus replication by approximately 1.3 log and 1.1 log respectively (Figure 3A and Figure S1A). In contrast, infected HIE that were pretreated with 25-HC in the presence of GCDCA/C2, required 5 µM 25-HC to suppress virus replication with a 5-fold decrease (Figure 3B and Figure S2B). Pretreatment of cells with 2.5 μM 24-HC and 27 27-HC, did not significantly reduce virus replication in the presence of GCDCA/C2 (data not shown).

**Figure 3:**
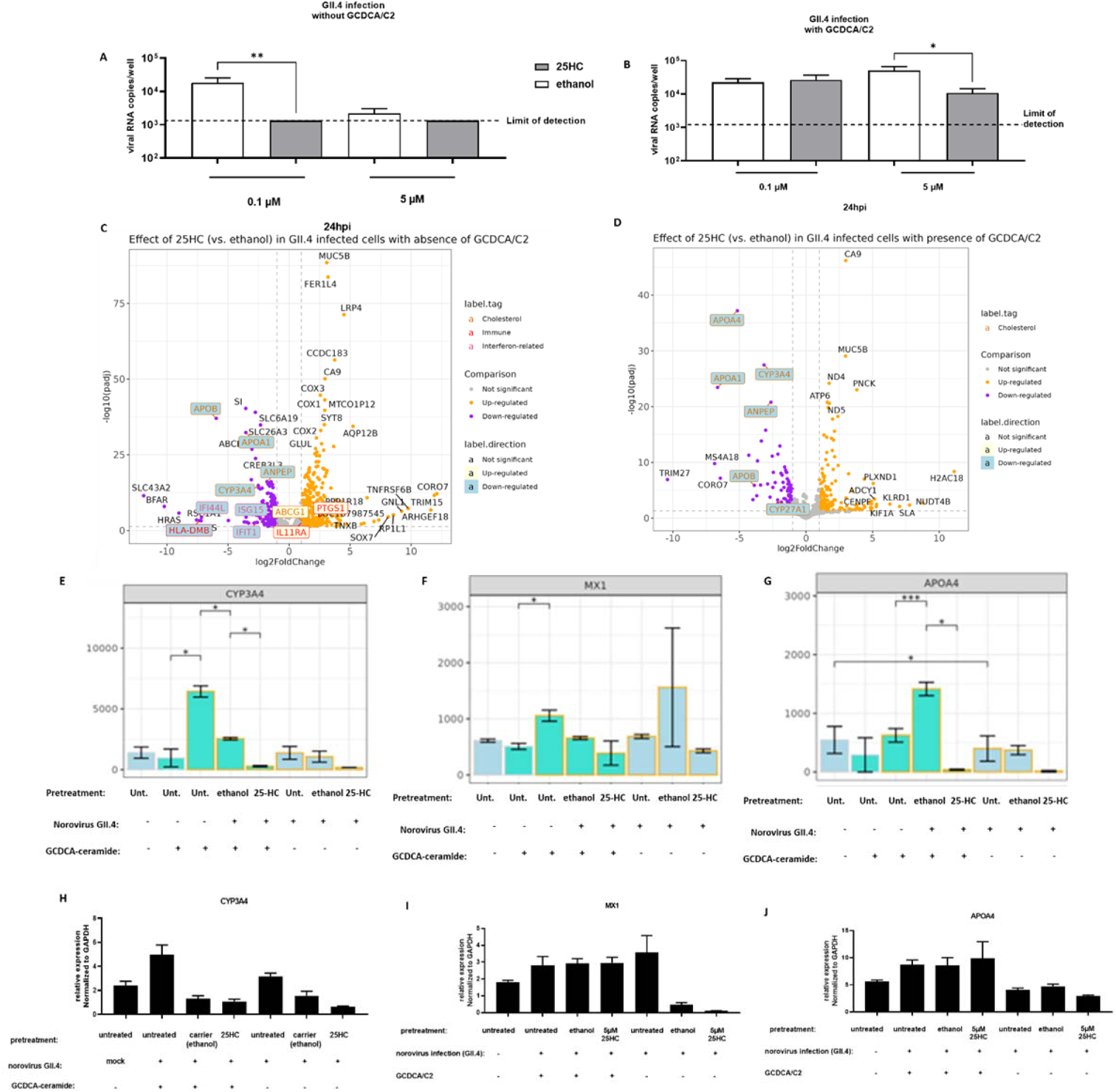
Gene expression in HIE monolayers pretreated with 0.1 μM 25-HC or equivalent dilution of ethanol (control) in for 24 hours and subsequently infected with human norovirus in media without GCDCA/C2 (A) or supplemented with 500μM GCDCA and 50μM ceramide (B). RNA was collected and analyzed by bulk RNA sequencing. Genes with absolute log2fold change higher than 1 and p-adjusted value < 0.05 were identified as the differentially expressed (DE) and are depicted in volcano plots (C) DE genes in cells infected in the absence of GCDCA/C2 of (D) presence of GCDCA/C2. Genes upregulated by 25-HC pre-treatment are shown in orange and genes downregulated by 25-HC pretreatment are shown in purple. (E-G) Normalized gene counts for CYP3A4, APOA4 and MX-1 as measured by bulk sequencing, (H-J) RT-PCR analysis of RNA isolated from cells treated and infected as indicated to detect expression of CYP3A4, APOA4 and MX-1. Gene expression was normalized to GAPDH. Two independent replicates per condition were analyzed by bulk-RNA sequencing. Data depicted is from one of two representative experiments with 3-4 replicates per condition.

To gain insights on the potential mechanism by which 25-HC restricts human norovirus and how 25-HC affects the transcriptional landscape during norovirus infection, we first determined the extent to which ethanol, in which 25-HC is dissolved, affected gene expression in human norovirus infected HIEs in the presence or absence of GCDCA/C2 (Figure S2). In the presence of GCDCA/C2 16 genes were differentially expressed between untreated cells and cells treated with the amount of ethanol equivalent to 0.1 μM 25-HC dilution (Figure S2) whereas in the presence of GCDCA/C2, 351 genes were differentially regulated as result of ethanol. Therefore, to rule out the impact of ethanol on gene expression, we used ethanol-treated HIE as baseline for gene expression data analyses for all 25-HC experiments.

In HIE pre-treated with 25-HC and infected with norovirus GII.4, 650 genes were DE in the absence of GCDCA/C2 (Figure 3C). Among the upregulated genes were mitochondrially encoded genes encoding subunits of the cytochrome C oxidase complex, and mitochondrial complex 1 as well as PTGS1 encoding cyclooxygenase 1. Other upregulated genes included those involved in the ATP metabolic process and cellular respiration including COX1, CO2 and COX3. Among downregulated genes were antiviral interferon-stimulated genes including IFI44L, IFIT1 MX-1 and ISG15 as well as genes also involved in cholesterol metabolism including CYP3A4 and CYP27A1 (Figure 3C).

Contrastingly, in the presence of GCDCA/C2, 235 genes were identified as DE genes (Figure 3D). Among upregulated genes were those associated with metabolic pathways such as mitochondrial genes ND4, ND5 and ATP6 as well as MUC5B and CA9 which are involved in mucus and acid production by intestinal cells. Among genes that were downregulated in response to 25-HC were APOA1 and APOA4, which are involved in cholesterol transport and associated with cholesterol metabolism along with CYP3A4 and CYP27A1genes.

Expression levels of genes that have been shown to be important in norovirus infection (MX-1), or 25-HC treatment (CYP3A4 and APOA1)(16, 43, 44) were confirmed by RT-qPCR (Figure 3E-J). CYP3A4 and MX1 were upregulated by norovirus infection but downregulated in 25-HC treated HIE and expression levels were even lower in infected HIE treated with 5 μM 25-HC in the absence of GCDCA/C2 (Figure 3E and F). Expression of APOA4 gene was not affected by norovirus, however, was decreased in response to 25-HC treatment in the presence or absence of GCDCA/C2. The gene expression for CYP3A4, MX1 and APOA4 was confirmed by RT-PCR (Figure 3H-J).

To gain additional insights into the cell signaling pathways affected by 25-HC pretreatment and subsequent norovirus infection in the absence and presence of GCDCA/C2, we performed a gene-network analysis using STRING (Figure 4). In the absence of GCDCA/C2, the pathways that were enriched by upregulated genes include viral transcription and viral gene expression, as well as protein trafficking to the endoplasmic reticulum. The pathways that were enriched by downregulated genes include the WNT signaling pathway, protein N-linked glycosylation pathway, fatty acid lipoprotein biosynthetic pathways type I interferon signaling as well as adaptive immune response (Figure 4A). Although fewer genes were differentially expressed overall in the presence of GCDCA/C2 (Figure 3D and S3), more pathways were identified that were enriched by the DE genes between 25-HC and ethanol treated RNA profiles. (Figure 4B). These pathways included those involved in the response to foreign substances (such as xenobiotic stimuli), lipid and lipoprotein biosynthesis, and cholesterol metabolism including cholesterol efflux.

**Figure 4:**
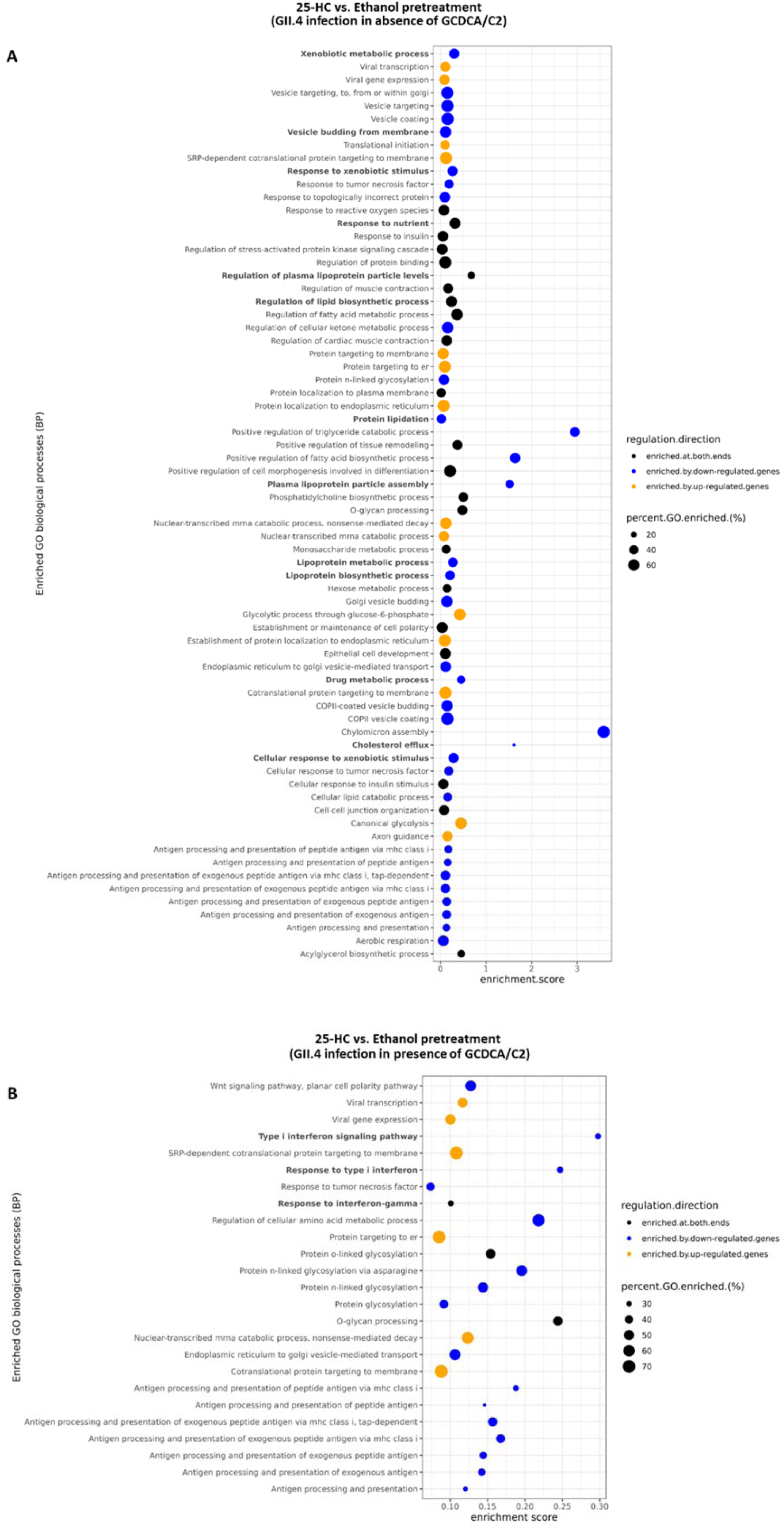
Pathway enrichment analysis showing pathways enriched by genes DE between the 25-HC pre-treated monolayers and the controls which were subsequently infected with norovirus in the absence (A) or presence (B) of GCDCA/C2. Cells were treated with 0.1μM 25-HC or equivalent dilution of ethanol (control) in for 24 hours and subsequently infected with human norovirus in the absence or presence of 500μM GCDCA and 50μM ceramide (C2) GCDCA/C2. RNA was collected and analyzed by bulk RNA sequencing. Pathways colored in orange are enriched predominantly by upregulated DE genes in this analysis and pathway colored in blue are enriched predominantly by down-regulated DE genes in this analysis. Some pathways were identified enriched by both up- and down-regulated genes and were colored in black; enrichment score reflects the degree of directionality (up- or down-regulated) of the genes enriching the pathway. The size of the dot is proportional to the percentage of the genes in the annotated GO pathways was enriched by the DE genes in this analysis. Pathway names highlighted in bold are pathways enriched by the genes of interest.

## Discussion

We report antiviral activity of 25-HC against human norovirus infected HIE. HIE that were treated with 25-HC prior to infection restricted virus replication in a dose-dependent manner. Similar findings have been reported for other viruses including rotavirus, severe acute respiratory syndrome coronavirus-2 (SARS-CoV-2), human immunodeficiency virus (HIV) and herpes simplex virus 1 (HSV1) (25, 29, 31, 45–47). We also found that GCDCA/C2 promote virus replication despite antiviral activity of 25-HC.

Previous data demonstrated that norovirus infections resulted in upregulation of antiviral interferon-stimulated genes in HIE (15, 16, 48, 49). In our study, antiviral genes such as IFIT1, MX-1 and ISG-15 were upregulated as well as CYP3A4, which is a gene that is involved cholesterol metabolism oxysterol biosynthesis, and production of 25-HC (50–52). We showed that 25-HC alters cellular gene expression in norovirus infected HIEs and downregulated cholesterol-related genes suggesting that 25-HC may affect cholesterol homeostasis. Suppression of genes related to cholesterol (CYP3A4 and APOA4) was partially alleviated by GCDCA/C2 although expression of these genes did not return to baseline level. In addition to suppression of cholesterol related genes, we found that 25-HC significantly upregulates mitochondrial genes related to cellular respiration.

A previous report using a Norwalk GI.1 replicon system demonstrated that Huh-7 cell genes related to cholesterol, lipid and metabolic pathways, cholesterol synthesis and intracellular esterification including HMG-CoA synthase, squalene epoxidase, LDLR-related protein 5 (LRP5) LRP12 LRP10 and ACAT2 were significantly downregulated in replicon-bearing cells, whereas SHP, LDLR class A domain containing 3 (LDLRAD3) and ACAT1 were significantly upregulated (53).

This study and our data underscore the importance of cholesterol/lipid pathways in norovirus infection. Investigation of mortality associated with a norovirus outbreak in a healthcare setting identified the use of cholesterol-lowering statins, as potential risk factors for acquiring symptomatic norovirus infection and excess mortality (54). Cholesterol synthesis and regulation of cholesterol by-products is complex, and further work needs to be done to determine the specific pathways important in human norovirus infection.

25-HC is a product of hydroxylation of cholesterol by cholesterol 25 hydroxylase, an enzyme encoded by the CH25H gene. 25-HC is a key regulatory molecule in cholesterol homeostasis which regulates intracellular cholesterol by regulating expression of enzymes involved in cholesterol biosynthesis as well as transport proteins involved in cholesterol uptake and efflux (55). CH25H is expressed at low basal levels in human cells but can be upregulated by interferon and is thus classified as an interferon-stimulated gene (ISG). As a result, 25-HC can be induced during virus infection and has been demonstrated to be broadly antiviral (56).

25-HC can also be produced via an alternative pathway via activity of cytochrome P450 3A4 (CYP3A4) an enzyme that is expressed in the liver and intestines (51). CYP3A4 has also been reported to be involved in bile metabolism (57). Thus, CYP3A4 may be involved in several pathways that are relevant to norovirus infection. We identified CYP3A4 as one of the genes that is significantly upregulated by human norovirus infection in untreated HIE cells, however; the role of CYP3A4 in norovirus infection has not been elucidated.

Bile acids support human norovirus replication in HIE. Although GII.4 viruses are capable of replicating without GCDCA and ceramide supplementation (11), it is possible that 25-HC suppresses cholesterol and expression of other genes necessary for downstream processes that lead to production of bile acids. Dysregulation of cholesterol/lipid homeostasis by 25-HC in the absence of GCDCA and C2 may explain why antiviral activity of 25-HC was only observed at higher concentrations (2.5 and 5 μM) in the presence of GCDCA and C2. One hypothesis is that human norovirus relies on cholesterol to support virus replication via production of bile salts. Although we have not established the exact mechanism by which 25-HC suppresses virus replication, we observed that 25-HC has profound effect on the oxysterol pathway and affects gene expression during infection and this may ultimately affect cholesterol metabolism or cellular distribution of cholesterol.

An unexpected finding in our study was that 25-HC treatment markedly reduced expression of antiviral ISGs. This was surprising, given the demonstrated antiviral activity of 25-HC against human norovirus. Our data suggest that the antiviral activity of 25-HC against human norovirus may be executed by a mechanism other than induction of ISGs. A recent report showed that 25-HC restricts Porcine Delta Coronavirus through suppression of TGFβ1 (58). 25-HC inhibits SARS-CoV-2 and other coronaviruses by depleting membrane cholesterol and/or by blocking membrane fusion (59) (30), while the cholesterol 25-hydroxylase (CH25H), which catalyze the production of 25-HC from cholesterol, suppresses porcine delta coronavirus infection by inhibiting viral entry (60). All these pathways may be worth exploring in future studies to understand the antiviral activity of 25-HC against human norovirus.

A limitation of the HIE system is that it does not account for other cell types that are found in the intestine that can sustain norovirus replication such as enteroendocrine epithelial cells(61). Specifically, the upregulation of 25-HC in response to interferon via the type I interferon receptor by dendritic cells and macrophages in response to type I interferon immune cells has been reported (62). However, we did not detect induction of CH25H by norovirus infection in HIE, despite other ISGs being induced. This does not rule-out the possibility that human norovirus does indeed induce CH25H in intestinal innate immune cells such as B cells, dendritic cells, macrophages innate lymphoid cells.

There are currently no approved antivirals to alleviate clinical norovirus symptoms, shorten duration of illness or reduce the burden of infection. The discovery that bile acids facilitate replication of human norovirus in HIE has revolutionized norovirus studies (11). Our study sheds light on the molecular complexity of this system and provides insights on potential targets for antiviral approaches as well as developing protocols that can improve virus yield. It also highlights the crosstalk between bile acids, innate immunity, and the antiviral response to norovirus (19, 63–65). Further studies may utilize these insights to optimize the HIE system and develop optimal cell culture conditions for increasing virus replication, serial passaging of the virus and eventually production of virus stocks, which has been a major limitation in the field.

---

**Figure S1:**
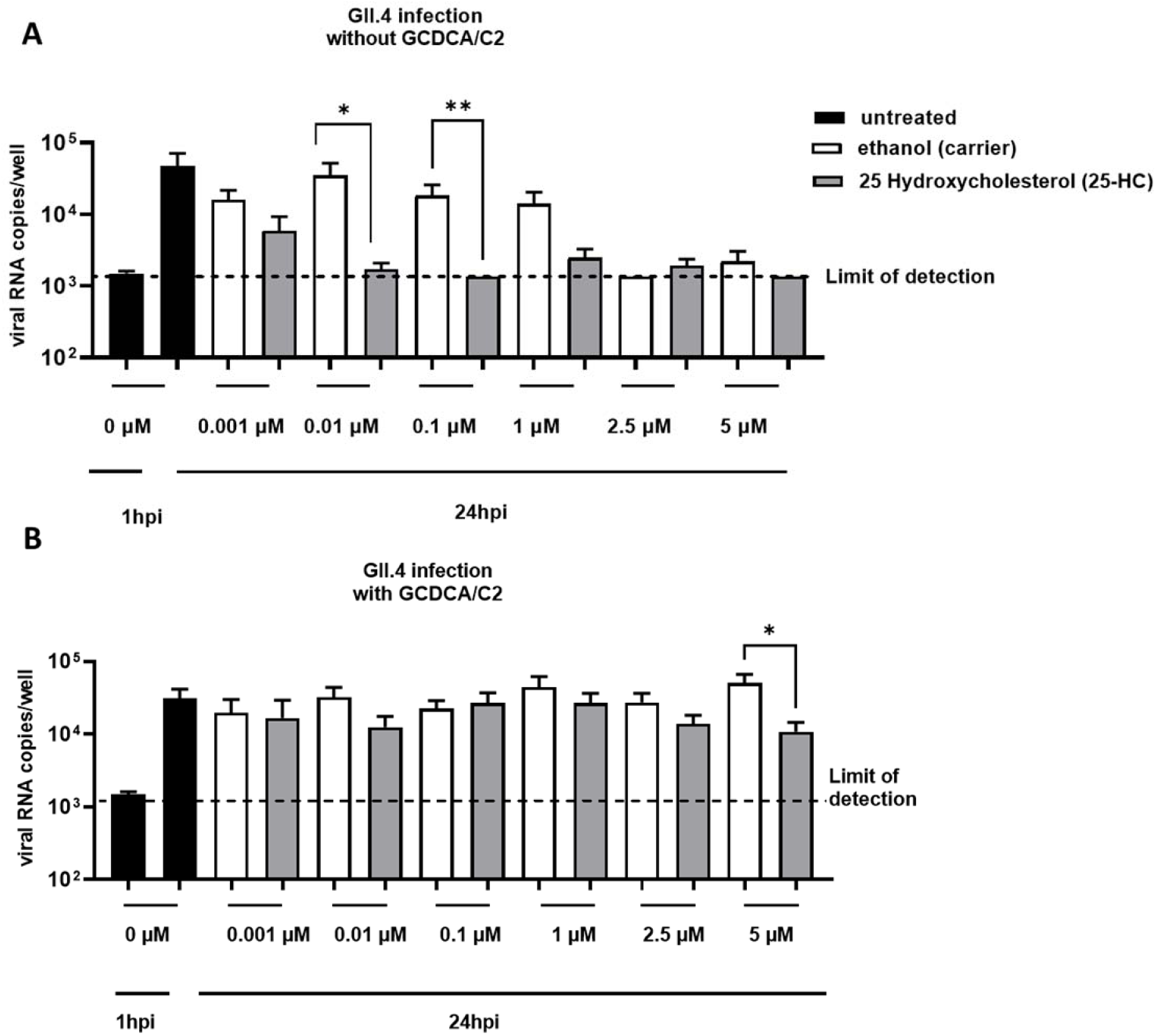
25-HC suppresses replication of human norovirus in human intestinal enteroid (HIE) cells. HIE monolayers were treated for 24 hours with 25-HC or equivalent dilution of the carrier (ethanol) in media then infected with norovirus GII.4 Sydney[P31] in the presence or absence of 500 μM GCDCA and 50 μM of ceramide (C2). (A) Quantification of norovirus RNA in cells infected in the absence of GCDCA/C2 at 24 hours post infection. (B) Quantification of norovirus RNA from cells infected in media in with GCDCA/C2. Data represents summary of 4 independent experiments with 3-4 independent replicates per condition in each experiment.

**Figure S2:**
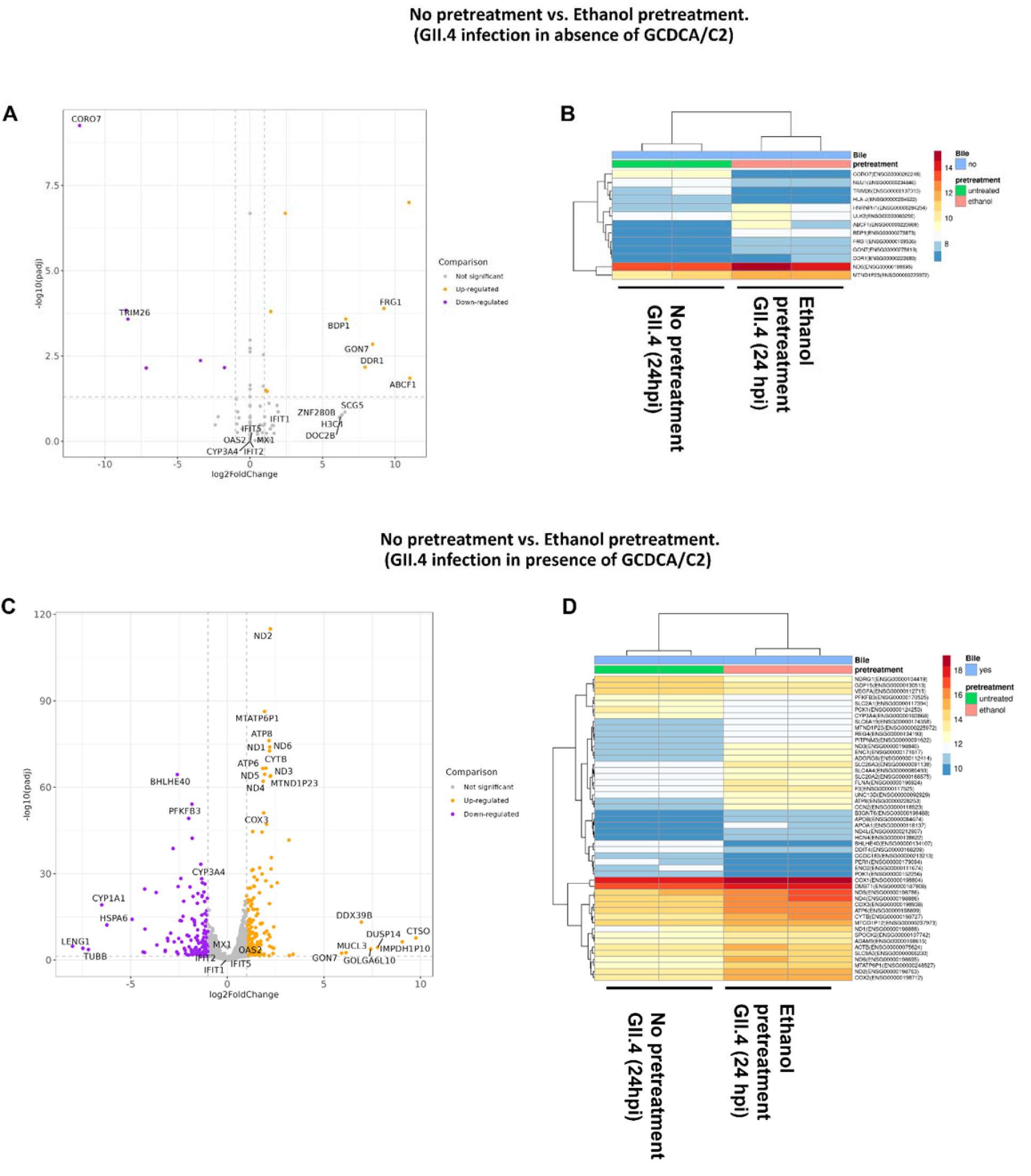
Effect of ethanol pretreatment on gene expression in HIE infected with human norovirus GII.4 in the absence or presence of GCDCA/C2. A. Volcano plot for DE genes of ethanol versus untreated with absence of GCDCA/C2. Genes upregulated by ethanol were colored in orange and genes downregulated by ethanol were colored in purple. B. Heat map. Relative gene expressions of all DE genes identified from the analysis. C. Volcano plot for DE genes of ethanol versus untreated with presence of GCDCA/C2 Genes upregulated by ethanol were colored in orange and genes downregulated by ethanol were colored in purple. D. Heat map. Relative gene expressions of the top50 DE genes identified from the analysis in C.

**Figure S3:**
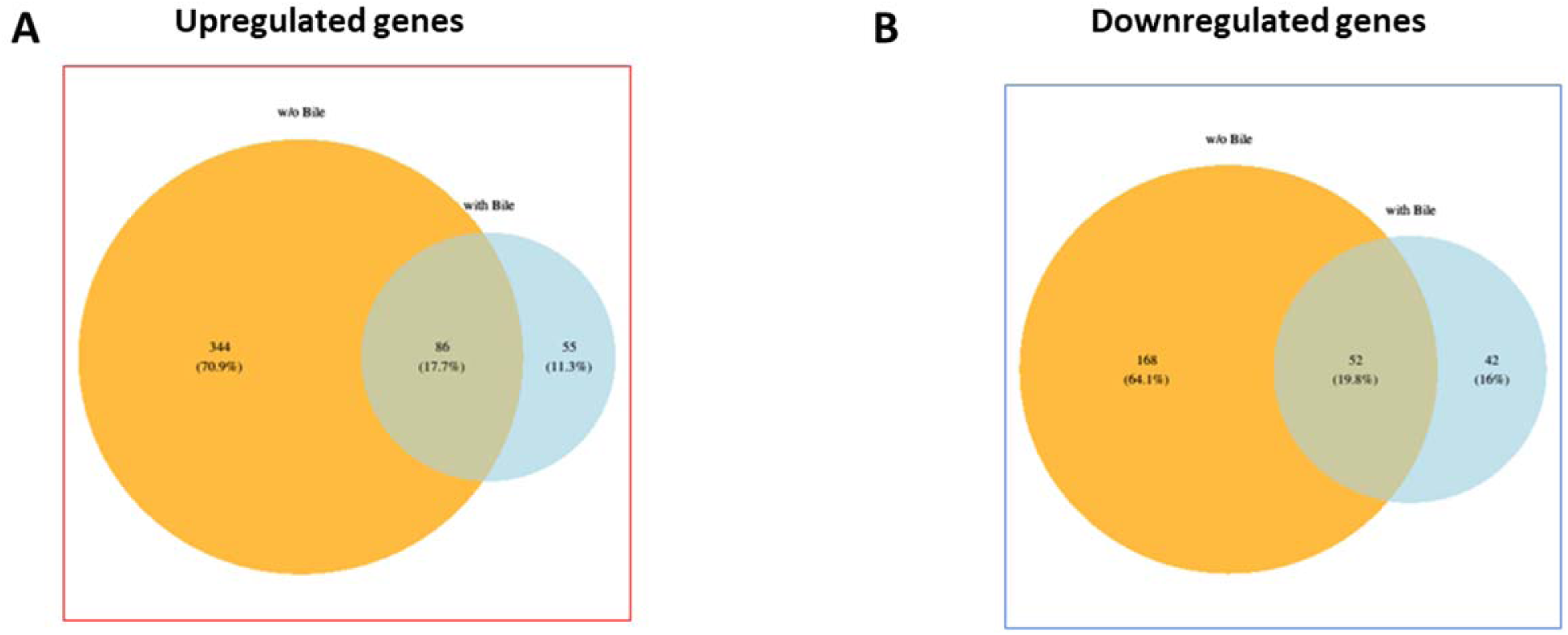
GCDCA/C2 affects 25-HC driven gene expression in norovirus infected HIE monolayers. Genes upregulated or downregulated in 25-HC treated HIE monolayers and subsequently infected norovirus in the presence of absence of GCDCA/C2 to determine overlapping and distinct genes. A) Venn diagram showing upregulated genes. B) Venn diagram showing downregulated genes.

## Supplemental information (SI)

**Table S1:**
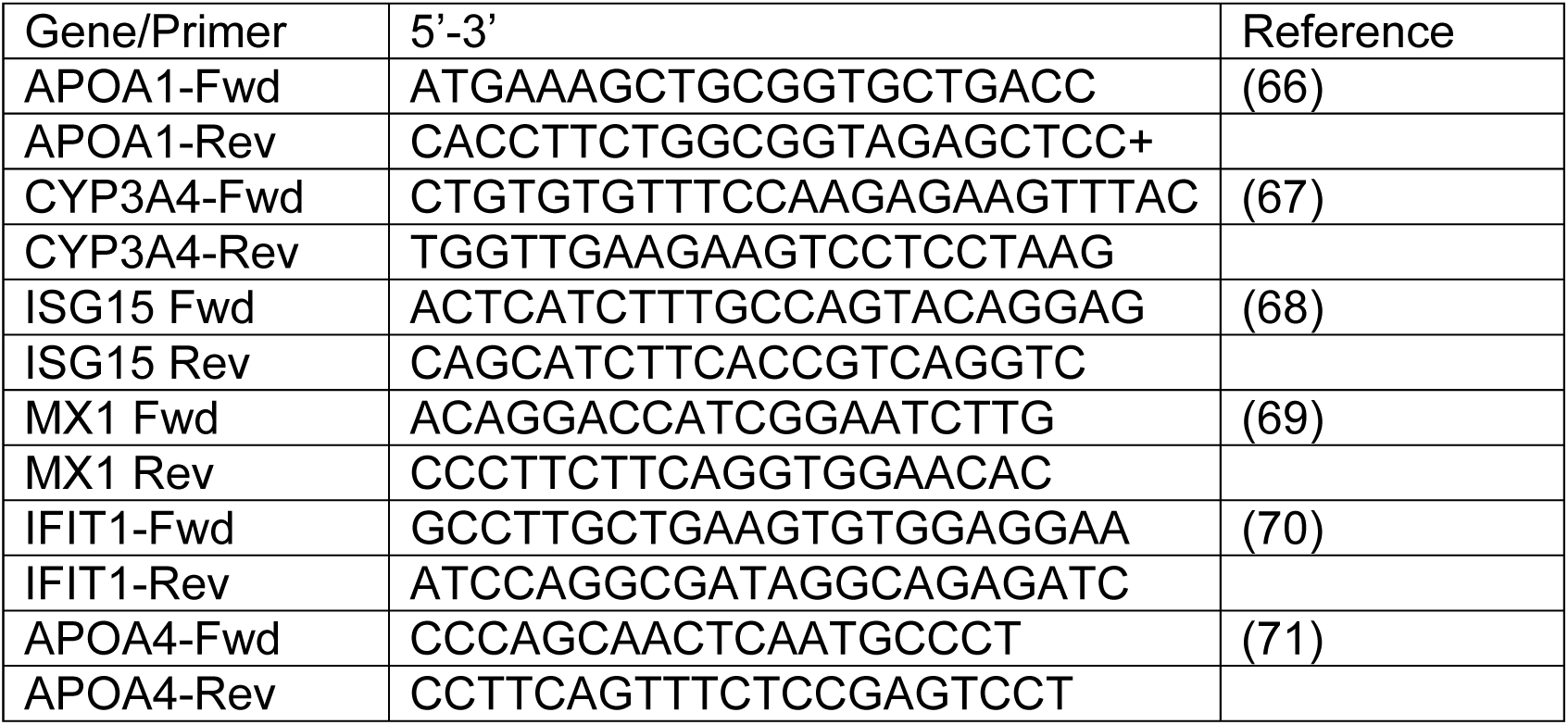
Oligonucleotide primers for amplifying select host genes

Genes upregulated by 25-HC in cells infected in the presence of bile and in the absence of bile (intersection)

MUC5B, FER1L4, LRP4, CCDC183, CA9, COX3, MTCO1P12, COX1, SYT8, AQP12B, COX2, GLUL, PSCA, MUC5AC, MUC2, ND5, CYTB, TFF1, ENO2, ND4, NA, ATP6, DUOXA2, NFATC4, CA4, NET1, CXCR4, ND4L, DAPK1, HLA-G, RAB13, FADS2, SLC6A8, DDIT4, TFF2, ND1, PFKFB3, GRIN2B, ELFN2, ABCA2, PTPRN2, TFF3, ND2, MTND2P28, BHLHE40, CHD3, LY6E, IFITM10, PPP1R1B, TRIB2, PPP1R3G, PNCK, TRAK2, ATP8, EGLN3, RAB3B, GAL3ST1, ANP32B, ANO1, BHLHE41, AK4, ALPK3, PLXND1, JAML, PDE9A, IGFBP6, TLCD2, NRP2, LYNX1, RP1L1, PPFIA4, FADS1, DUX4, ZNF469, CEMIP, TMC4, NOTCH3, SPDEF, FLG, MUC6, MUC16, MUCL3, MTATP6P1, ND3

Genes upregulated only shown in absence of bile

HLA-G, SH3PXD2A, ECM1, GOLGA8A, IGFBP2, DUOX2, COL27A1, CTSD, SPAG4, VEGFA, LMNTD2, KIFC2, SHC2, SULT1C2, COL5A3, NDRG1, S100A4, CAPN8, LPIN3, BIRC7, CLIP2, ZNF395, CDC42BPG, ADAM8, CDH23, SRRM2, ARHGAP26, CORO7, FGD5, TRIM15, LDC1P, PLEKHA4, SLCO4A1, CCDC88B, GPI, PPP1R18, CAPN12, INSC, DNM1, ZBTB7C, SLC34A2, FREM1, DVL1, RASSF7, SLC2A1, CLDN23, ATP2A3, PDK1, CCN4, NLGN2, ADM, PLCXD1, CLK3, TRERF1, TBC1D3D, CENPB, AKR1C2, PFKP, AJM1, TRIM31, ATG16L2, MAGED4B, MLPH, RASSF3, AKNA, TBC1D3E, WASH8P, LENG8, SCART1, SORL1, KIF1C, NFKBIA, TNFRSF6B, NPAS3, PHKA2, ST8SIA6, COL6A1, ARHGEF18, TNFAIP3, C22orf46, CLASRP, FGF11, DOT1L, PPARD, EFNA1, ALOX5, PALM3, GNL1, SPIRE2, RBBP8NL, TRIP10, WNK2, LOC613038, TFRC, SLC29A4, PLCH2, NA, COQ8A, C1orf115, PGK1, SNX8, LMTK3, RASA4B, NBPF19, ZSWIM9, RNF215, UGT2B15, AKR1C1, PDE10A, MST1L, COL1A1, STC2, KANK2, SPACA6, HILPDA, CSPG4, CCHCR1, SPIDR, ZBTB47, CGNL1, TSEN2, HNRNPUL2-BSCL2, TLE2, UGT2B17, TMEM141, NBPF20, S100A6, SOX7, TNNT1, EFEMP2, SLC9A3R2, OSBPL5, PITPNM3, SYNE3, GPR37L1, FRY, KAT2A, ABI3BP, NOXO1, PTGS1, HSPA1A, FCGBP, S100A14, STC1, DYRK1B, SH3BP1, SLC25A45, NPIPB3, ANKFN1, MXRA8, RNA5SP202, GNE, SNCG, KRI1, HLA-J, OTOP3, AP2A2, NOL12, WSCD1, C15orf48, ING1, CELSR3, SOCS3, HSF4, NECAB3, CACNA1E, N4BP3, CA12, MYH10, BCO1, AGAP4, MIF, PRDM10, HIF3A, TNFSF9, ADAMTS13, COL28A1, TNXB, TINCR, NOMO1, MXD3, SHF, ALDOC, RNF152, RAB26, GPX2, LIG1, TNNI2, RNF183, ST3GAL4, SELENBP1, RPL23A, LOC107987545, IGIP, UAP1L1, ABCG1, EFNA3, CD70, EHD2, ITGB2, SYT7, SLFN11, SCARF1, PDE4C, NLRP1, C3, P3H3, MTND4P12, NDRG2, FOXA2, SPOCD1, C20orf96, ADAMTS14, IL11RA, EGFL7, SLC6A12, MMP28, RAB31, SPTSSB, C17orf78, GRIN2A, GRK3, H1-3, DEF6, LOC441242, ATP8B3, TNS2, RAPGEF4, TTLL10, TPTEP2-CSNK1E, ICAM5, RAB37, TRAPPC5, PLIN5, MROH7, RILP, C1QL1, PBX4, DIRAS2, MSH5, SRCIN1, MPV17L, PIGZ, IL17REL, ADCY4, CPNE7, CDX1, GDF11, FREM2, GAS2L3, SRRM3, NMB, MASP1, EBF2, EVL, RELT, ZNF467, PAXBP1, RRAGD, KCNH2, MAFB, ATP5MF-PTCD1, PCDH9, DDAH2, DUOXA1, CDH4, GABRA2, CLIC3, KRT42P, FSCN2, SFTA2, GALNT18, PAK6, LSAMP, ABLIM3, FAM177B, GFAP, CCDC146, RBP3, GOLGA6L9, FOSL1, CALB1, PPARGC1A, JAK3, TRIM7, CRABP2, JPH1, ARMH1, KIAA1549L, LMNB1, SERPINA4, LGI4, CCDC80, BAIAP3, GLP1R, CSKMT, CACNA1C, AQP12A, RECQL4, MTRNR2L2, S1PR4, GRAP, PCOLCE, PRSS2, SETBP1, POLN, MKI67, POU4F1, LOC645433, TLE6, RGS19, CACNA2D3, EIF4EBP3, TAF4B, IRX2, EEF2K, KLK1, ZNF251

Genes upregulated genes only in the presence of bile:

MUCL3, H2AC18, PROM2, PKM, MUC2, SCD, RDH11, MYEOV, STARD10, ADCY1, MUC6, NKD1, CAPN9, CHD6, CLIC1, NA, DNAH11, RGPD2, KLRD1, PROX1, LOXL4, KIF1A, NUDT4B, PRR36, MYO16, PAG1, STARD9, TF, SLA, CENPF, TGM4, ANGPTL4, NOMO3, GRM2, NCF4, DUX4L26, SLC12A5, LOXL1, RARG, OTULINL, KNOP1, DERL3, CARMIL2, RBFOX3, LINC02693, C14orf132, RPL23AP21, OR10H1, UBXN11, CEACAM7, CELSR1, IKZF2, CEP152

Genes downregulated by 25-HC in cells infected in the presence of bile and in the absence of bile (intersection)

CYP3A4, APOA1, ANPEP, CREB3L3, RBP2, CDHR2, GGT1, SLC6A19, SLC43A2, AQP3, SLC3A2, SMIM5, ACE, SLC46A1, VTN, HLA-DRB1, MXRA5, MUC13, MAF, ALDOB, MTTP, ACSL5, SUSD2, APOB, SLC26A3, RARRES1, CYBRD1, ACE2, FOLH1, TRPV6, ISX, GDA, REG1A, MS4A8, HCN4, GLS, PLS1, HMOX1, BTNL3, NABP1, ABCB1, PID1, ABCG5, GPRIN1, GCNT4, GPCPD1, PAQR7, GK, HSD17B11, LPCAT3, ADA, B3GNT6

Genes down-regulated only in absence of bile:

SI, ENC1, AASS, RAB11FIP3, MGAM, MYO1B, SLC43A2, SLC20A2, ODC1, TRPV6, SFRP5, SLC4A8, ALDH1A3, SPOCK2, MROH6, YIPF4, SLC7A8, CLDN1, GALC, CLCN5, C1QTNF5, POFUT1, BFAR, CEACAM20, TMEM236, AP2B1, PPP2CB, TSPAN13, SAMD9L, FAM3C, MYL9, POLR1A, DPP4, PLAGL2, VTCN1, HRAS, CMPK2, LIPA, DTX3L, SMIM31, TM2D2, CIDEB, CHST15, ANKRD29, CLDN2, MEP1B, NPPB, MMP24, NIPAL1, ENTPD7, FRMD1, HMGN2, HLA-DRB1, HSPB8, GPD1, DDC, SLC35G1, OAS3, MN1, IFI44L, TMEM87B, KRT23, LEPROT, ACOT7, ELOVL5, CASP1, SLC7A9, ZMPSTE24, CADM1, TGFB2, SDSL, SLC46A3, FAM20A, HPGD, ISG15, FAM20C, NA, FSTL1, ABCG8, RSC1A1, ZMYND15, CFTR, MVB12B, CDKN1C, HLA-DMB, NTS, DDX52, NUDT9, XPNPEP2, EIF2S1, IRF8, SNN, LAP3, JKAMP, PIBF1, PTGIS, HPS5, IFI30, C8G, SNX11, DAB2, ALPI, ASAH2, RNF43, AGPAT4, BMP8A, MARVELD2, LAMP3, CPSF3, XAF1, IFIT1, PLA2G12B, UFSP1, TMEM151A, RIMKLB, TMEM192, AQP10, FBXL17, COL4A1, PSMC1P1, REG1B, C2, EPSTI1, GGT1, SIRPA, OIT3, FAM47E-STBD1, SLC13A2, SRPRB, CDK7, IFIT3, PNPT1, CLCN4, PCSK1N, ARHGEF26, CPQ, NEU4, HLA-DRA, SULT1B1, GBP4, PALMD, COA7, CD37, CCDC170, BST1, DDX58, TTC13, SLC2A9, SP110, HLA-DRB3, ZNF850, ZNF268, PKIG, CNOT3, SBSPON, PSMB3, COQ2, SMIM11A, C2orf88, CMTM3, ZNF678, ZNF441, ADH1C, USP18

Genes down-regulated genes only in the presence of bile:

APOA4, MS4A18, THBS1, CORO7, TRIM27, SELENOP, GLYCTK, SLC9A3, GPA33, DGAT1, CD74, CES2, CDHR5, UGT1A1, RECQL5, ID3, ETS2, SECTM1, PRR5L, A1CF, SLC15A1, CYP27A1, MEP1A, VIPR1, GSDME, HLA-DRA, SERPINA1, SMAD7, TM4SF20, OAT, SLC5A1, CIITA, MYO1A, GUCY2C, NEURL1B, PRAP1, SCTR, PCK2, GREB1L, EPHX2, FABP2, MOGAT3

## References

1. Ahmed SM, Hall AJ, Robinson AE, Verhoef L, Premkumar P, Parashar UD, Koopmans M, Lopman BA. 2014. Global prevalence of norovirus in cases of gastroenteritis: a systematic review and meta-analysis. Lancet Infect Dis 14:725–730.

2. Hall AJ, Lopman BA, Payne DC, Patel MM, Gastanaduy PA, Vinje J, Parashar UD. 2013. Norovirus disease in the United States. Emerg Infect Dis 19:1198–205.

3. Liao Y, Hong X, Wu A, Jiang Y, Liang Y, Gao J, Xue L, Kou X. 2021. Global prevalence of norovirus in cases of acute gastroenteritis from 1997 to 2021: An updated systematic review and meta-analysis. Microb Pathog 161:105259.

4. Kirk MD, Pires SM, Black RE, Caipo M, Crump JA, Devleesschauwer B, Dopfer D, Fazil A, Fischer-Walker CL, Hald T, Hall AJ, Keddy KH, Lake RJ, Lanata CF, Torgerson PR, Havelaar AH, Angulo FJ. 2015. World Health Organization Estimates of the Global and Regional Disease Burden of 22 Foodborne Bacterial, Protozoal, and Viral Diseases, 2010: A Data Synthesis. PLoS Med 12:e1001921.

5. Lucero Y, Vidal R, O’Ryan GM. 2018. Norovirus vaccines under development. Vaccine 36:5435–5441.

6. Hallowell BD, Parashar UD, Hall AJ. 2019. Epidemiologic challenges in norovirus vaccine development. Hum Vaccin Immunother 15:1279–1283.

7. Mattison CP, Cardemil CV, Hall AJ. 2018. Progress on norovirus vaccine research: public health considerations and future directions. Expert Rev Vaccines 17:773–784.

8. Costantini V, Morantz EK, Browne H, Ettayebi K, Zeng XL, Atmar RL, Estes MK, Vinje J. 2018. Human Norovirus Replication in Human Intestinal Enteroids as Model to Evaluate Virus Inactivation. Emerg Infect Dis 24:1453–1464.

9. Ramani S, Crawford SE, Blutt SE, Estes MK. 2018. Human organoid cultures: transformative new tools for human virus studies. Curr Opin Virol 29:79–86.

10. Ettayebi K, Tenge VR, Cortes-Penfield NW, Crawford SE, Neill FH, Zeng XL, Yu X, Ayyar BV, Burrin D, Ramani S, Atmar RL, Estes MK. 2021. New Insights and Enhanced Human Norovirus Cultivation in Human Intestinal Enteroids. mSphere 6.

11. Ettayebi K, Crawford SE, Murakami K, Broughman JR, Karandikar U, Tenge VR, Neill FH, Blutt SE, Zeng XL, Qu L, Kou B, Opekun AR, Burrin D, Graham DY, Ramani S, Atmar RL, Estes MK. 2016. Replication of human noroviruses in stem cell-derived human enteroids. Science 353:1387–1393.

12. Mboko WP, Chhabra P, Valcarce MD, Costantini V, Vinje J. 2022. Advances in understanding of the innate immune response to human norovirus infection using organoid models. J Gen Virol 103.

13. Nolan LS, Baldridge MT. 2022. Advances in understanding interferon-mediated immune responses to enteric viruses in intestinal organoids. Front Immunol 13:943334.

14. Crawford SE, Ramani S, Blutt SE, Estes MK. 2021. Organoids to Dissect Gastrointestinal Virus-Host Interactions: What Have We Learned? Viruses 13.

15. Jahun AS, Goodfellow IG. 2021. Interferon responses to norovirus infections: current and future perspectives. J Gen Virol 102.

16. Hosmillo M, Chaudhry Y, Nayak K, Sorgeloos F, Koo BK, Merenda A, Lillestol R, Drumright L, Zilbauer M, Goodfellow I. 2020. Norovirus Replication in Human Intestinal Epithelial Cells Is Restricted by the Interferon-Induced JAK/STAT Signaling Pathway and RNA Polymerase II-Mediated Transcriptional Responses. mBio 11.

17. Arthur SE, Sorgeloos F, Hosmillo M, Goodfellow IG. 2019. Epigenetic Suppression of Interferon Lambda Receptor Expression Leads to Enhanced Human Norovirus Replication In Vitro. mBio 10.

18. Murakami K, Tenge VR, Karandikar UC, Lin SC, Ramani S, Ettayebi K, Crawford SE, Zeng XL, Neill FH, Ayyar BV, Katayama K, Graham DY, Bieberich E, Atmar RL, Estes MK. 2020. Bile acids and ceramide overcome the entry restriction for GII.3 human norovirus replication in human intestinal enteroids. Proc Natl Acad Sci U S A 117:1700–1710.

19. Tenge VR, Murakami K, Salmen W, Lin SC, Crawford SE, Neill FH, Prasad BVV, Atmar RL, Estes MK. 2021. Bile Goes Viral. Viruses 13.

20. Willinger T. 2019. Oxysterols in intestinal immunity and inflammation. J Intern Med 285:367–380.

21. Lembo D, Cagno V, Civra A, Poli G. 2016. Oxysterols: An emerging class of broad spectrum antiviral effectors. Mol Aspects Med 49:23–30.

22. Abrams ME, Johnson KA, Perelman SS, Zhang LS, Endapally S, Mar KB, Thompson BM, McDonald JG, Schoggins JW, Radhakrishnan A, Alto NM. 2020. Oxysterols provide innate immunity to bacterial infection by mobilizing cell surface accessible cholesterol. Nat Microbiol 5:929–942.

23. Brown AJ, Jessup W. 2009. Oxysterols: Sources, cellular storage and metabolism, and new insights into their roles in cholesterol homeostasis. Mol Aspects Med 30:111–22.

24. Pandak WM, Kakiyama G. 2019. The acidic pathway of bile acid synthesis: Not just an alternative pathway(). Liver Res 3:88–98.

25. Civra A, Francese R, Gamba P, Testa G, Cagno V, Poli G, Lembo D. 2018. 25-Hydroxycholesterol and 27-hydroxycholesterol inhibit human rotavirus infection by sequestering viral particles into late endosomes. Redox Biol 19:318–330.

26. Li C, Deng YQ, Wang S, Ma F, Aliyari R, Huang XY, Zhang NN, Watanabe M, Dong HL, Liu P, Li XF, Ye Q, Tian M, Hong S, Fan J, Zhao H, Li L, Vishlaghi N, Buth JE, Au C, Liu Y, Lu N, Du P, Qin FX, Zhang B, Gong D, Dai X, Sun R, Novitch BG, Xu Z, Qin CF, Cheng G. 2017. 25-Hydroxycholesterol Protects Host against Zika Virus Infection and Its Associated Microcephaly in a Mouse Model. Immunity 46:446–456.

27. Liu SY, Aliyari R, Chikere K, Li G, Marsden MD, Smith JK, Pernet O, Guo H, Nusbaum R, Zack JA, Freiberg AN, Su L, Lee B, Cheng G. 2013. Interferon-inducible cholesterol-25-hydroxylase broadly inhibits viral entry by production of 25-hydroxycholesterol. Immunity 38:92–105.

28. Wu C, Zhao J, Li R, Feng F, He Y, Li Y, Huang R, Li G, Yang H, Cheng G, Chen L, Ma F, Li P, Sun C. 2021. Modulation of Antiviral Immunity and Therapeutic Efficacy by 25-Hydroxycholesterol in Chronically SIV-Infected, ART-Treated Rhesus Macaques. Virol Sin 36:1197–1209.

29. Serquina AKP, Tagawa T, Oh D, Mahesh G, Ziegelbauer JM. 2021. 25-Hydroxycholesterol Inhibits Kaposi’s Sarcoma Herpesvirus and Epstein-Barr Virus Infections and Activates Inflammatory Cytokine Responses. mBio 12:e0290721.

30. Zang R, Case JB, Yutuc E, Ma X, Shen S, Gomez Castro MF, Liu Z, Zeng Q, Zhao H, Son J, Rothlauf PW, Kreutzberger AJB, Hou G, Zhang H, Bose S, Wang X, Vahey MD, Mani K, Griffiths WJ, Kirchhausen T, Fremont DH, Guo H, Diwan A, Wang Y, Diamond MS, Whelan SPJ, Ding S. 2020. Cholesterol 25-hydroxylase suppresses SARS-CoV-2 replication by blocking membrane fusion. Proc Natl Acad Sci U S A 117:32105–32113.

31. Shawli GT, Adeyemi OO, Stonehouse NJ, Herod MR. 2019. The Oxysterol 25-Hydroxycholesterol Inhibits Replication of Murine Norovirus. Viruses 11.

32. Cannon JL, Barclay L, Collins NR, Wikswo ME, Castro CJ, Magana LC, Gregoricus N, Marine RL, Chhabra P, Vinje J. 2017. Genetic and Epidemiologic Trends of Norovirus Outbreaks in the United States from 2013 to 2016 Demonstrated Emergence of Novel GII.4 Recombinant Viruses. J Clin Microbiol 55:2208–2221.

33. Cannon JL, Barclay L, Collins NR, Wikswo ME, Castro CJ, Magana LC, Gregoricus N, Marine RL, Chhabra P, Vinje J. 2019. Correction for Cannon et al., “Genetic and Epidemiologic Trends of Norovirus Outbreaks in the United States from 2013 to 2016 Demonstrated Emergence of Novel GII.4 Recombinant Viruses”. J Clin Microbiol 57.

34. Chen S, Zhou Y, Chen Y, Gu J. 2018. fastp: an ultra-fast all-in-one FASTQ preprocessor. Bioinformatics 34:i884–i890.

35. Patro R, Duggal G, Love MI, Irizarry RA, Kingsford C. 2017. Salmon provides fast and bias-aware quantification of transcript expression. Nat Methods 14:417–419.

36. HochbergYosef BYa. 1995. Controlling the False Discovery Rate: A Practical and Powerful Approach to Multiple Testing. Journal of the Royal Statistical Society Series B (Methodological) Vol. 57:289–300.

37. (ed). 2016. ggplot2: Elegant Graphics for Data Analysis. Accessed

38. Szklarczyk D, Gable AL, Lyon D, Junge A, Wyder S, Huerta-Cepas J, Simonovic M, Doncheva NT, Morris JH, Bork P, Jensen LJ, Mering CV. 2019. STRING v11: protein-protein association networks with increased coverage, supporting functional discovery in genome-wide experimental datasets. Nucleic Acids Res 47:D607–D613.

39. Cao QT, Aguiar JA, Tremblay BJ, Abbas N, Tiessen N, Revill S, Makhdami N, Ayoub A, Cox G, Ask K, Doxey AC, Hirota JA. 2020. ABCF1 Regulates dsDNA-induced Immune Responses in Human Airway Epithelial Cells. Front Cell Infect Microbiol 10:487.

40. Lee MN, Roy M, Ong SE, Mertins P, Villani AC, Li W, Dotiwala F, Sen J, Doench JG, Orzalli MH, Kramnik I, Knipe DM, Lieberman J, Carr SA, Hacohen N. 2013. Identification of regulators of the innate immune response to cytosolic DNA and retroviral infection by an integrative approach. Nat Immunol 14:179–85.

41. Ma Z, Moore R, Xu X, Barber GN. 2013. DDX24 negatively regulates cytosolic RNA-mediated innate immune signaling. PLoS Pathog 9:e1003721.

42. Serfecz JC, Hong Y, Gay LA, Shekhar R, Turner PC, Renne R. 2022. DExD/H Box Helicases DDX24 and DDX49 Inhibit Reactivation of Kaposi’s Sarcoma Associated Herpesvirus by Interacting with Viral mRNAs. Viruses 14.

43. Mears HV, Emmott E, Chaudhry Y, Hosmillo M, Goodfellow IG, Sweeney TR. 2019. Ifit1 regulates norovirus infection and enhances the interferon response in murine macrophage-like cells. Wellcome Open Res 4:82.

44. Rodriguez MR, Monte K, Thackray LB, Lenschow DJ. 2014. ISG15 functions as an interferon-mediated antiviral effector early in the murine norovirus life cycle. J Virol 88:9277–86.

45. Civra A, Cagno V, Donalisio M, Biasi F, Leonarduzzi G, Poli G, Lembo D. 2014. Inhibition of pathogenic non-enveloped viruses by 25-hydroxycholesterol and 27-hydroxycholesterol. Sci Rep 4:7487.

46. Dong H, Zhou L, Ge X, Guo X, Han J, Yang H. 2018. Antiviral effect of 25-hydroxycholesterol against porcine reproductive and respiratory syndrome virus in vitro. Antivir Ther 23:395–404.

47. Tricarico PM, Caracciolo I, Gratton R, D’Agaro P, Crovella S. 2019. 25-hydroxycholesterol reduces inflammation, viral load and cell death in ZIKV-infected U-87 MG glial cell line. Inflammopharmacology 27:621–625.

48. Lin L, Han J, Yan T, Li L, Li J, Ao Y, Duan Z, Hou Y. 2019. Replication and transcriptionomic analysis of human noroviruses in human intestinal enteroids. Am J Transl Res 11:3365–3374.

49. Cieza RJ, Golob JL, Colacino JA, Wobus CE. 2021. Comparative Analysis of Public RNA-Sequencing Data from Human Intestinal Enteroid (HIEs) Infected with Enteric RNA Viruses Identifies Universal and Virus-Specific Epithelial Responses. Viruses 13.

50. Diczfalusy U, Bjorkhem I. 2011. Still another activity by the highly promiscuous enzyme CYP3A4: 25-hydroxylation of cholesterol. J Lipid Res 52:1447–9.

51. Honda A, Miyazaki T, Ikegami T, Iwamoto J, Maeda T, Hirayama T, Saito Y, Teramoto T, Matsuzaki Y. 2011. Cholesterol 25-hydroxylation activity of CYP3A. J Lipid Res 52:1509–16.

52. Wang Y, Yutuc E, Griffiths WJ. 2021. Cholesterol metabolism pathways - are the intermediates more important than the products? FEBS J 288:3727–3745.

53. Chang KO. 2009. Role of cholesterol pathways in norovirus replication. J Virol 83:8587–95.

54. Rondy M, Koopmans M, Rotsaert C, Van Loon T, Beljaars B, Van Dijk G, Siebenga J, Svraka S, Rossen JW, Teunis P, Van Pelt W, Verhoef L. 2011. Norovirus disease associated with excess mortality and use of statins: a retrospective cohort study of an outbreak following a pilgrimage to Lourdes. Epidemiol Infect 139:453–63.

55. Cao Q, Liu Z, Xiong Y, Zhong Z, Ye Q. 2020. Multiple Roles of 25-Hydroxycholesterol in Lipid Metabolism, Antivirus Process, Inflammatory Response, and Cell Survival. Oxid Med Cell Longev 2020:8893305.

56. Mao S, Ren J, Xu Y, Lin J, Pan C, Meng Y, Xu N. 2022. Studies in the antiviral molecular mechanisms of 25-hydroxycholesterol: Disturbing cholesterol homeostasis and post-translational modification of proteins. Eur J Pharmacol 926:175033.

57. Bodin K, Lindbom U, Diczfalusy U. 2005. Novel pathways of bile acid metabolism involving CYP3A4. Biochim Biophys Acta 1687:84–93.

58. Zhang J, Yang G, Wang X, Zhu Y, Wang J. 2022. 25-Hydroxycholesterol Mediates Cholesterol Metabolism to Restrict Porcine Deltacoronavirus Infection via Suppression of Transforming Growth Factor beta1. Microbiol Spectr doi:10.1128/spectrum.02198-22:e0219822.

59. Wang S, Li W, Hui H, Tiwari SK, Zhang Q, Croker BA, Rawlings S, Smith D, Carlin AF, Rana TM. 2020. Cholesterol 25-Hydroxylase inhibits SARS-CoV-2 and other coronaviruses by depleting membrane cholesterol. EMBO J 39:e106057.

60. Ke W, Wu X, Fang P, Zhou Y, Fang L, Xiao S. 2021. Cholesterol 25-hydroxylase suppresses porcine deltacoronavirus infection by inhibiting viral entry. Virus Res 295:198306.

61. Green KY, Kaufman SS, Nagata BM, Chaimongkol N, Kim DY, Levenson EA, Tin CM, Yardley AB, Johnson JA, Barletta ABF, Khan KM, Yazigi NA, Subramanian S, Moturi SR, Fishbein TM, Moore IN, Sosnovtsev SV. 2020. Human norovirus targets enteroendocrine epithelial cells in the small intestine. Nat Commun 11:2759.

62. Park K, Scott AL. 2010. Cholesterol 25-hydroxylase production by dendritic cells and macrophages is regulated by type I interferons. J Leukoc Biol 88:1081–7.

63. Fiorucci S, Biagioli M, Zampella A, Distrutti E. 2018. Bile Acids Activated Receptors Regulate Innate Immunity. Front Immunol 9:1853.

64. Podevin P, Rosmorduc O, Conti F, Calmus Y, Meier PJ, Poupon R. 1999. Bile acids modulate the interferon signalling pathway. Hepatology 29:1840–7.

65. Wang J, Flavell RA, Li HB. 2019. Antiviral immunity: a link to bile acids. Cell Res 29:177–178.

66. Tsezou A, Iliopoulos D, Malizos KN, Simopoulou T. 2010. Impaired expression of genes regulating cholesterol efflux in human osteoarthritic chondrocytes. J Orthop Res 28:1033–9.

67. Murray GI, McFadyen MC, Mitchell RT, Cheung YL, Kerr AC, Melvin WT. 1999. Cytochrome P450 CYP3A in human renal cell cancer. Br J Cancer 79:1836–42.

68. Enosi Tuipulotu D, Netzler NE, Lun JH, Mackenzie JM, White PA. 2018. TLR7 Agonists Display Potent Antiviral Effects against Norovirus Infection via Innate Stimulation. Antimicrob Agents Chemother 62.

69. Zhang S, Kodys K, Li K, Szabo G. 2013. Human type 2 myeloid dendritic cells produce interferon-lambda and amplify interferon-alpha in response to hepatitis C virus infection. Gastroenterology 144:414–425 e7.

70. Yi X, Cheng X. 2021. Understanding Competitive Endogenous RNA Network Mechanism in Type 1 Diabetes Mellitus Using Computational and Bioinformatics Approaches. Diabetes Metab Syndr Obes 14:3865–3945.

71. Fan Y, Gao J, Li Y, Chen X, Zhang T, You W, Xue Y, Shen C. 2021. The Variants at APOA1 and APOA4 Contribute to the Susceptibility of Schizophrenia With Inhibiting mRNA Expression in Peripheral Blood Leukocytes. Front Mol Biosci 8:785445.

